# Rheological behavior of Pluronic/Pluronic diacrylate hydrogels used for bacteria encapsulation

**DOI:** 10.1101/2023.03.29.534703

**Authors:** Shardul Bhusari, Maxi Hoffmann, Petra Herbeck-Engel, Shrikrishnan Sankaran, Manfred Wilhelm, Aránzazu del Campo

## Abstract

Pluronic (Plu) hydrogels containing Pluronic diacrylate (PluDA) have become popular matrices to encapsulate bacteria in engineered living materials. For this purpose, 30 wt% Plu/PluDA hydrogels with variable fraction of covalently crosslinkable PluDA in the hydrogel composition are used. The degree of covalent crosslinking and the consequent different mechanical properties of the hydrogels have been shown to affect bacteria growth, but a systematic investigation of the mechanical response of the hydrogels is still missing. Here we study the rheological response of 30 wt.% Plu/PluDA hydrogels with increasing PluDA fraction between 0 and 1. We quantify the range of viscoelastic properties that can be covered in this system by varying in the PluDA fraction. We present stress relaxation and creep-recovery experiments, and analyze the variation of the critical yield strain/stress, relaxation and recovery parameters of Plu/PluDA hydrogels as function of the covalent crosslinking degree using the Burgers and Weilbull models. We expect this study to help users of Plu/PluDA hydrogels to estimate the mechanical properties of their systems, and eventually to correlate them with the behaviour of bacteria in future Plu/PluDA devices of similar composition.

## 1 Introduction

Soft hydrogel networks with combined physical (dynamic) and covalent (permanent) crosslinks are interesting materials for the encapsulation and controlled expansion of cells.^1,2^ The inherent reorganization capability of the dynamic crosslinks allows cells to deform the surrounding material as they proliferate and accommodate daughter cells. In parallel, the elasticity of the network imposes an increasing compressive force on the cell population as it grows and controls its size.^3,4^ Hydrogels formed by mixtures of Pluronic (PEO_x_-PPO_y_-PEO_x_) and Pluronic diacrylate (PluDA) have been successfully used to embed and regulate the growth of bacteria and yeast in engineered living materials.^1,5,6,7,8,9^ Despite the positive results and the application potential of the developed systems, a systematic analysis of the mechanical properties of these hydrogels is still missing.

Physical Pluronic hydrogels are formed by aggregation of the Pluronic micelles into clusters through the interactions between the PEO coronas (**Scheme 1**). Above the transition temperature, the micellar clusters grow and build a 3D network.^10,11^ For the specific Pluronic F127 (PEO_106_-PPO_70_-PEO_106_), frequently used in bacteira encapsulation and also for the experiments in this article, the critical micellar concentration (cmc) is at 0.725 wt. % in water at 25 °C ^12–14^, and the sol-gel transition occurs at polymer concentrations > 5 wt. % and temperatures above 14 °C. The assembly of Pluronic F127 chains in 3D networks has been studied by scattering methods like dynamic light scattering (DLS) and small angle neutron scattering (SANS).^14,15^ The order of the micelles in the clusters depends on external conditions (i.e., temperature, salt concentration, shear forces)^15,16^ and ranges from perfect face-centered cubic (FCC) or hexagonally close packed (HCP) structures to random stackings. As associative physical networks, Pluronic hydrogels show elastic properties in the quiescent state and undergo yielding, i.e. fracture and flow, when the stress surpasses the interparticle forces.^17,18^ The shear thinning behavior and the thermosensitivity makes Pluronic gels interesting from a biomedical perspective.^19^ It allows them to be easily mixed with cells or other payloads and processed using shear forces.^20–22^

**Scheme 1:**
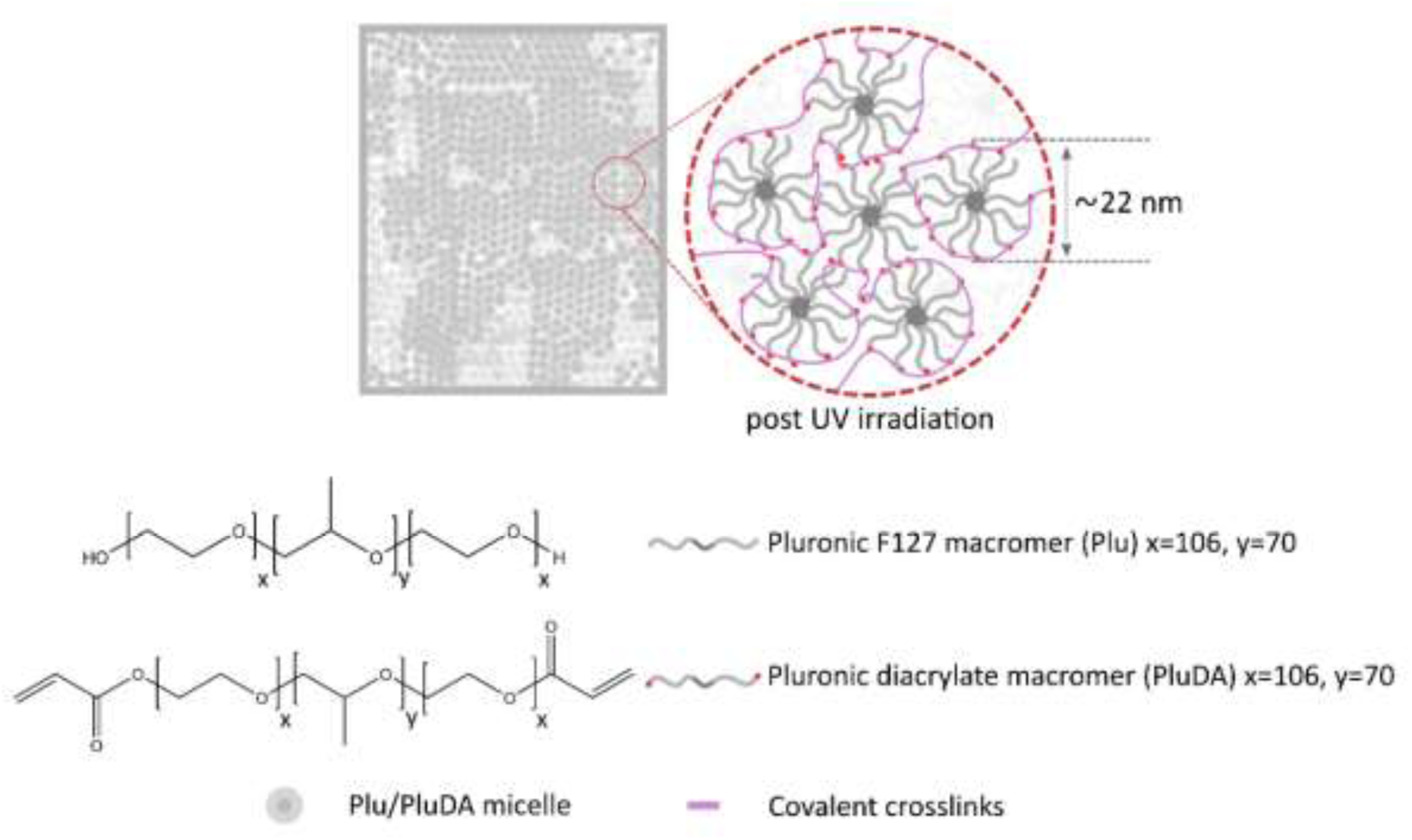
Micellar structure of the Plu/PluDA hydrogel representing the organization of the PEOx-PPOy-PEOx triblock copolymer chains above the sol-gel transition temperature. The micelles have a diameter of ca. 22 nm and build aggregates that form a network. with the inter- and intra-micellar covalent crosslinks between PluDA chains.

Physical Pluronic hydrogels swell and dissociate into individual micelles when immersed in water. The introduction of polymerizable acrylate functionalities as end-groups of the Pluronic chains provides the possibility to stabilize the micellar hydrogel via the formation of permanent, covalent bonds between the micelles.^7,11^ Covalent crosslinking affects the mechanical behavior, specially the elasticity as quantified via G’ of the hydrogel.^1^ By varying the PluDA fraction in Plu/PluDA hydrogels the density of chemical crosslinks in the hydrogel and the resulting mechanical properties can be tuned. For example, a 30 wt. % Plu/PluDA hydrogels show increasing elastic response (G’) from 17.8 ± 1.4 to 42.9 ± 1.9 kPa and decreasing non-elastic response (from creep-recovery curves) when the PluDA fraction in the mixture increased from 0 to 100%.^1^

Here we present a detailed study of the time-dependent rheological behavior of Plu/PluDA hydrogels as a function of PluDA fraction. We use 30 wt. % Plu/PluDA hydrogels as used in preceding reports for the fabrication of engineered living materials by different groups.^6,7,23,24^

## 2 Materials and Methods

### 2.1 Sample preparation

Pluronic diacrylate (PluDA) was synthesized by reaction of Pluronic F127 (Plu) with acryloyl chloride in the presence of triethylamine according to a reported protocol.^11^ Acrylation degrees of 70% were typically obtained.

Plu/PluDA hydrogels were prepared by mixing stock solutions of Plu and PluDA polymers as previously reported.^1^ The resulting hydrogels were named DA X, with X being the fraction of PluDA in the Plu/PluDA mixture, between 0 and 100 %. 30 wt. % Plu and PluDA stock solutions (named DA0 and DA100, respectively) were prepared in milliQ water and contained 0.2% w/v Irgacure 2959 (Sigma-Aldrich Co.) as photoinitiator. Solutions were stored at 4 °C. DA 25, DA 50 and DA 75 hydrogels were prepared by mixing DA0 and DA100 stock solutions in the following ratios-3:1 (DA 25), 1:1 (DA 50) and 1:3 (DA 75). After mixing, the DA X hydrogels were allowed to form at room temperature for 10 minutes.^25^ For the photoinitiated crosslinking, hydrogels were exposed to UV light (365 nm, 6 mW/cm^2^) using a OmniCure Series 1500 lamp for 60 s through a UV transparent bottom plate for rheology.

### 2.2 Raman spectroscopy of Plu/PluDA powders and DA 0-100 hydrogels

Raman investigations were carried out at ambient conditions on a LabRAM HR Evolution HORIBA Jobin Yvon A Raman microscope (Longmujeau, France) using a 633 nm He–Ne laser (Melles Griot, IDEX Optics and Photonics, Albuquerque, NM, USA) equipped with 1800 lines per 1 mm grating.

### 2.3 Rheological measurements

The rheological properties were measured with a stress-controlled rheometer (DHR 3, TA Instruments) using a parallel plate geometry. A 20 mm Peltier plate/ UV transparent plate was used as bottom plate and a smooth stainless steel 12 mm disk was used as top plate. The rheometer was equipped with a UV Source (OmniCure, Series 1500, 365 nm, 6 mW/cm^2^) for illumination of the hydrogel samples in between the rheometer plates. Experiments were performed at room temperature (22-23°C). To avoid drying of the sample by evaporation during testing a solvent trap was used and the sample was sealed with silicone oil.

The 30 wt. % DA X hydrogels were prepared by pipetting 35 µL of a freshly prepared 30 wt. % DA X precursors solution on the rheometer plate, and allowing the gel to form between the plates (diameter 12 mm, gap 300 µm) during 10 min. The polymerization of the acrylate groups was initiated by exposure to 365 nm (6 mW/cm^2^) through the UV transparent bottom plate. Strain sweeps were conducted from 0.001% to 1000% at a frequency of 1 Hz. In the strain amplitude sweep curves, a line was fit to the plateau region (linear) and the drop-off region (non-linear). The intersection is taken as the critical strain value (*γ*_y_).^26,27^ Fluidization strain (*γ*_F_) was taken as the value at the intersection of G’ and G’’, i.e., at G’ = G’’).^27^ The corresponding stress values were used as critical stress (τ_y_) and fluidization stress (τ_F_) values. The experimental conditions for the rheological experiments were taken from our previous work.^1^

### 2.4 Stress relaxation measurements

In the stress relaxation experiment, strains of 0.5, 1, 2, 5, 10, 15 and 30 % were applied to the same sample (DA 0, 50 and 100, n=3) for 300 s with a strain raise time of 0.2 s. To avoid cumulative stress buildup and shear history effects, the strain was reversed (negative strain, equivalent to the positive strain) after each run to reach rheometer stress = 0 Pa, and the next run was started after 10 min equilibration time.

In another experiment, a constant strain of 1% was applied to the sample (strain raise time 0.01 s) and the stress was monitored for 300 s. Each experiment was repeated three times.

The percentage of relaxation was defined as the drop in the relaxation modulus from the start (30 ms) to the end (300 s) of the experiment normalized by the starting relaxation modulus.^29^

The relaxation curves were fitted to a linear combination of two stretched exponential or Kohlrausch–Williams–Watts (KWW) functions,^28^

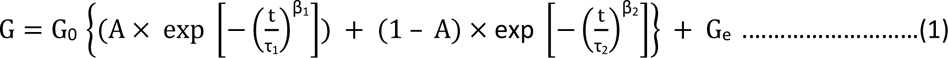

where G_0_ is the relaxation modulus linearly extrapolated to zero time, τ_1_ and τ_2_ are the viscoelastic relaxation times for the two processes (0 < τ), β_1_ and β_2_ are stretching exponents for the two processes (0 < β ≤ 1) that reflect the width of the relaxation time distribution.

The parameter A is the fractional contribution of the fast relaxation to the whole relaxation process (0 < A) and G_e_ is the equilibrium relaxation shear modulus.

### 2.5 Creep recovery measurements

A shear stress of 100 Pa was applied for 180 s to the hydrogel sample in the rheometer. The shear strain was monitored during this time (creep phase), and for a further 180 s after removal of the shear stress (recovery phase). Creep ringing phenomenon was observed at short time scales (2 s) in the creep and recovery experiments.^29^

Creep deformation, γ _creep_, was fitted to a four parameter Burgers model:^30–32^

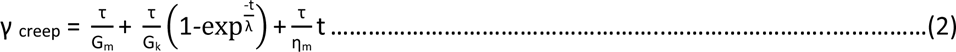

where τ is the applied shear stress (100 Pa), t denotes the time after loading, G_m_ denotes the shear modulus (or spring constant) of the spring, η_m_ denotes the viscosity of the dashpot in the Maxwell element, G_k_ denotes the shear modulus (or spring constant) and η_k_ denotes the viscosity of the dashpot in the Kelvin element, and λ is the retardation time (η_k_/G_k_) needed to achieve 63.2% of the total deformation in the Kelvin unit. The non-linear curve fit function of the OriginPro 9.1 software was used for the fitting and four parameters (G_m_, G_k_, η_m_ and τ) were defined. Creep ringing phenomenon was observed at short time scales. Therefore only data at t > 2 s were considered in the fitting process.^29^

Recovery data was fitted to a Weibull distribution function.^30,33^

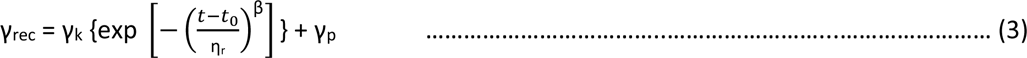

where where γ_rec_ denotes the deformation after the instantaneous strain recovery, γ_k_ denotes the delayed viscoelastic strain recovery (Kelvin-Voigt element), η_r_ is the characteristic life parameter, t_0_ is an induction time and β the shape factor. The stress is removed at time t_0_ (180 s) and γ_p_ is the permanent irreversible strain. Ringing phenomenon was observed again at short time scales as in creep data. Therefore, t_0_ = 182 s was considered for the fitting.

### 2.6 Statistical Analyses

Non-linear curve fit function of the OriginPro 9.1 software was used for the fitting of the rheological curves. Concrete parameters used for the different types of analyses performed have been described in the sections and figure captions of the related experiments.

## 3 Results

The 30 wt. % Plu/PluDA hydrogels were prepared by mixing defined volumes of 30 wt. % solutions of Plu and PluDA polymers.^1^ Hydrogels were named DA X, with X being the fraction of PluDA in the Plu/PluDA mixture, between 0 and 100 % (**Table 1**). In the following we describe first the rheological behavior of the physical DA X hydrogels, and of the physical and covalently crosslinked DA X hydrogels.

**Table 1:** Composition and mechanical parameters of DA X hydrogels. Note that the end-group functionalization degree of PluDA component is 70% and, therefore, the real fraction of covalent fixed chains differs for each composition and reaches 70% as maximum value in DA 100.

| Hydrogel name (DA X) | Plu:PluDA ratio (mol %) | Ratio of diacrylated endgroups<sup>1</sup> (%) | Shear storage modulus<sup>2</sup> G' (kPa) | Strain amplitude at yield point<sup>2</sup> $\gamma_y$ (%) | Strain amplitude at fluidization<sup>2</sup> $\gamma_f$ (%) | Critical stress<sup>3</sup> $\tau_y$ (kPa) | Fluidization stress<sup>4</sup> $\tau_f$ (kPa) |
| --- | --- | --- | --- | --- | --- | --- | --- |
| DA 0 | 100 : 0 | 0 | 15.3 ± 1.9 | 1.8 ± 0.6 | 5.5 ± 1.3 | 0.2 ± 0.0 | 0.2 ± 0.0 |
| DA 25 | 75 : 25 | 17.5 | 21.2 ± 2.1 | 6.0 ± 0.4 | 24.1 ± 2.8 | 0.9 ± 0.0 | 1.7 ± 0.0 |
| DA 50 | 50 : 50 | 35 | 25.1 ± 4.2 | 9.5 ± 1.2 | 46.4 ± 7.3 | 1.7 ± 0.3 | 3.9 ± 0.2 |
| DA 75 | 25 : 75 | 52.5 | 37.1 ± 0.7 | 22.8 ± 6.9 | 61.1 ± 21.4 | 5.2 ± 0.8 | 6.7 ± 0.6 |
| DA 100 | 0 : 100 | 70 | 47.5 ± 2.9 | 30.0 ± 14.7 | 65.0 ± 31.3 | 8.3 ± 3.1 | 9.9 ± 0.9 |

<sup>1</sup> calculated taking into account 70% end-group functionalization of PluDA
<sup>2</sup> from strain amplitude sweep, values at $\gamma_0 = 0.1$ % and frequency = 1 Hz (**Figure 1a**)
<sup>3</sup> corresponding stress values of $\gamma_y$
<sup>4</sup> corresponding stress values of $\gamma_f$

### 3.1 Rheological behavior of physical Plu and PluDA hydrogels

A 30 wt. % solution of Pluronic F127 (Plu) in water forms physical hydrogels at temperature >15.5°C.^1^ At room temperature (22°C) and shear strain amplitudes (𝛾_0_) below 1 %, the Plu (= DA 0) hydrogel behaves as a linear response viscoelastic solid (**Figure 1a**) with a strain amplitude independent shear modulus G’ of 15.3 ± 1.6 kPa (**Figure 1b**) at ω/2π = 1 Hz. At higher strain amplitudes (γ_0_ > 1%), Plu hydrogels show strain-induced yielding and gradual fluidization (**Figure 1a**). This behavior is characteristic of colloidal or granular hydrogels,^34^ in which particles aggregate through reversible interparticle interactions to form an interconnected, percolated 3D network structure that yields when the applied stress exceeds the interparticle attractive forces.^26^ The critical strain for yielding mainly depends on the particle content and the strength of the interparticle interactions including interparticle (electrostatic, hydrogen bonding, van der Waals, steric, depletion etc.) and hydrodynamic forces.^35^

**Figure 1.**
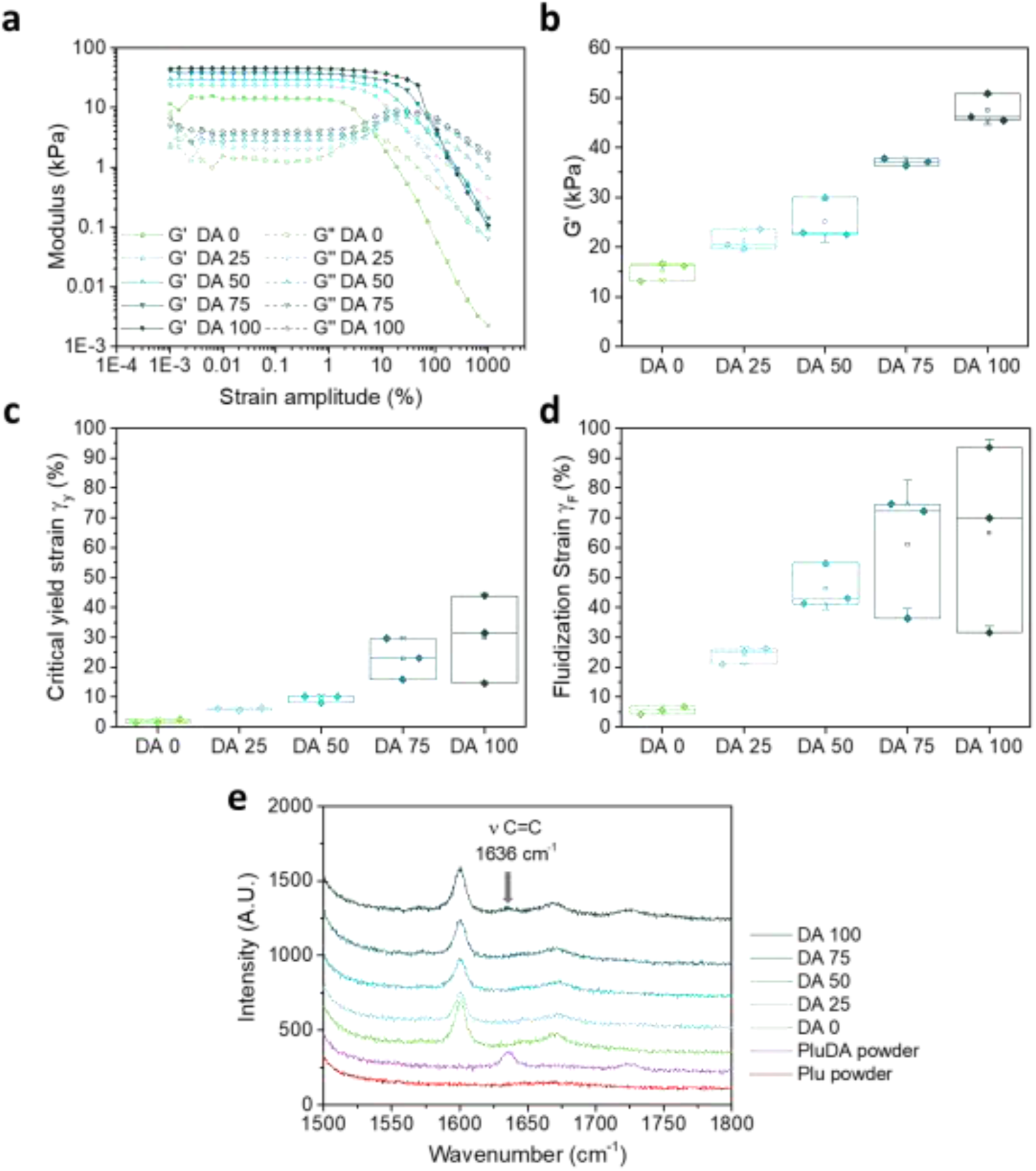
**a**) Representative strain amplitude sweeps of DA X hydrogels measured at a frequency of 1 Hz; **b)** G’ values of DA X hydrogels from (a) at 0.1 % strain amplitude at a frequency of 1 Hz; **c)** Values of strain amplitude at critical yield point, γy, and **d)** strain amplitude at fluidization point, γF, of DA X hydrogels as function of X obtained from the strain amplitude sweep experiment in **a**), (N = 3, box represents 25 and 75 percentile values and whiskers indicate standard deviation); **e)** Raman spectra (vertically shifted for clarity) of Plu and PluDA powders and DA X hydrogels. PluDA powder shows the C=C stretching mode (ν) at 1636 cm^-1^. The band at 1600 cm^-1^ in DA X hdyrogels corresponds to the -OH bending mode of water molecules, which is absent in Plu and PluDA powders.

The 30 wt. % solutions of diacrylated Pluronic F127 (PluDA, 70 % end-group functionalization) also form physical hydrogels at a slightly lower temperature of 14°C.^1^ The substitution of the terminal –OH groups by less polar acrylic functionalities in the outer surface of the PEO shell of Pluronic micelles enhances micellar aggregation and gel formation.^11^ The resulting physical PluDA hydrogel shows a similar G’ in the linear viscoelastic region (**Figure S1a**) and a two-fold higher critical strain amplitude for yielding in the strain sweep experiment (**Figure S1b**) compared to the Plu hydrogel, confirming stronger inter-micellar interactions in the gels with acrylate end-groups.

The strain sweep curves of the physical Plu and PluDA hydrogels show an overshoot in G’’ at intermediate strain amplitudes between the linear viscoelastic and the fluidization regions. This overshoot is referred to as the Payne effect and is related to yielding as a gradual transition that occurs while the deformation increases.^36^ Strain-dependent breakdown of the internal structure, length scale-dependent rearrangements or forced stress relaxation are believed to contribute to the overshoot of G’’ as a function of applied strain amplitude.^36^

### 3.2 Rheological behavior of physically vs. physically and covalently crosslinked PluDA hydrogel

Covalently crosslinked PluDA (=DA100) hydrogels were prepared from 30 wt. % PluDA solutions containing 0.2 wt. % Irgacure 2959. The solution was left to form a physical gel between the rheometer plates, and it was covalently crosslinked subsequently by exposure to 365 nm light through the UV transparent bottom plate. The degree of conversion of the acrylate groups after polymerization was evaluated by Raman spectroscopy. The characteristic band for the stretching of the C=C bond, observed at 1635 cm^-1^ in the spectrum of PluDA powder, diminished in Plu powder and in the PluDA hydrogel after UV irradiation (**Figure 1e**). This indicates nearly full conversion of the acrylate groups in the photo-crosslinking step.

The polymerization of the acrylate groups influenced the rheological response of the PluDA hydrogel. The covalently crosslinked hydrogel showed a broader linear viscoelastic range than the exclusively physically crosslinked PluDA, with strain amplitude independent moduli up to 𝛾_0_ = 10% (**Figure S1a**). At higher strain amplitudes, a continuous drop in G’ was observed. This strain induced yielding and gradual fluidization suggests that the micellar network in the covalent crosslinked PluDA hydrogel can undergo major reorganization and stress dissipation and stress release in spite of 70% of the endgroups of the polymeric chains being covalently crosslinked. This behavior can be understood considering that acrylate groups at the PluDA chain terminals can form inter- and intra-micellar crosslinks during the photoinduced polymerization reaction.^11^ Micellar clusters which are covalently crosslinked are expected to contribute to the elasticity of the hydrogel and be the reason for the higher G’ and higher critical yielding strain of the covalently crosslinked PluDA hydrogel compared to the physically crosslinked PluDA hydrogel (30 ± 14.7 v*s.* 4.2 ± 0.8 %, **Figure 1c, S1b**). Intra-micellar crosslinks stabilize the micelles and form loops in the chains. These crosslinks do not interfere with the inherent ability of physical micellar aggregates to flow above the critical strain. Note that the average residence time of a Pluronic molecule in a physical micellar aggregate has been estimated to be several hours.^37^ Therefore, crosslinking during photopolymerization (1 min time scale) can mainly occur between chain ends located in close proximity. The observed strain-induced yielding and fluidization (**Figure 1d**) above a critical yield stress in our PluDA hydrogels suggests that a significant fraction of the covalent crosslinks are intra-micellar. This has been previously suggested by other authors based on gel permeation chromatography (GPC) and light scattering analysis of the sol phase after crosslinking and cooling below the transition temperature.^11^

### 3.3 Rheological behavior of covalently crosslinked Plu/PluDA hydrogels with variable PluDA content

We analysed the rheological response of Plu/PluDA covalent hydrogels as a function of the degree of covalent crosslinking. Near full conversion after the photo-crosslinking step was confirmed by Raman spectroscopy (**Figure 1e**).

Covalently crosslinked DA X hydrogels showed an increasing trend of G’, *γ*_y_ and *γ*_F_ values from Plu (=DA 0) to PluDA (=DA 100) hydrogels. The strain amplitude sweeps showed both a linear viscoelastic and a fluidization region. The storage modulus G’ at 0.1 % strain amplitude increased from 15.3 ± 1.8 to 47.5 ± 2.9 kPa as the PluDA content increased from 0 to 100 % (**Figure 1c**). The critical strain for yielding at 1 Hz, *γ_y_*, also increased from 1.7 ± 0.4 to 29.7 ± 15.2 % (**Figure 1d**, **Table 1**) from DA 0 to DA 100. The strain amplitude needed to reach the fluidization point (crossover of G’’ and G’, as opposite to gelation point), *γ_F_,* increased from 5.5 ± 1.2 to 65.0 ± 31.3 % (**Figure 1e**) and the corresponding stress values at the fluidization point ranged from 0.2 ± 0.0 to 9.9 ± 0.9 kPa (**Figure S2, Table 1**). The increase in the density of covalent links between neighboring micelles (inter-micellar) with increasing X hinders the viscous deformation of the hydrogel and provides higher mechanical stability and elasticity under the applied strain.

### 3.4 Stress relaxation of DA X hydrogels

Figure 2a-c shows the stress induced in DA 0, DA 50 and DA 100 hydrogels during a stress relaxation experiment with step-strain increase from 0.5 to 30 % (300 s) and strain raise time of 200 ms. The stress relaxation curves normalized by the maximum stress values attained during the 200 ms strain raise are also shown. In physical DA 0 hydrogels, the maximum induced stress increased from 78 ± 17 to 231 ± 55 Pa with increasing applied strain from 0.5 to 5 %, and it plateaued for larger shear deformations (Figure 2d). In contrast, the maximum induced stress in covalently crosslinked DA 100 increased linearly with the strain up to 15 % and reached values up to 5708 ± 246 Pa (Figure 2d), denoting a primarily elastic behavior within this strain range. DA 50 showed an intermediate behavior. These differences are in agreement with the observations from the strain sweep experiment (Figure 1c), where *γ*_y_ of physical DA 0 hydrogel was 1.8 ± 0.6% while that for covalently crosslinked DA 100 hydrogel was 30.0 ± 14.7 %.

**Figure 2:**
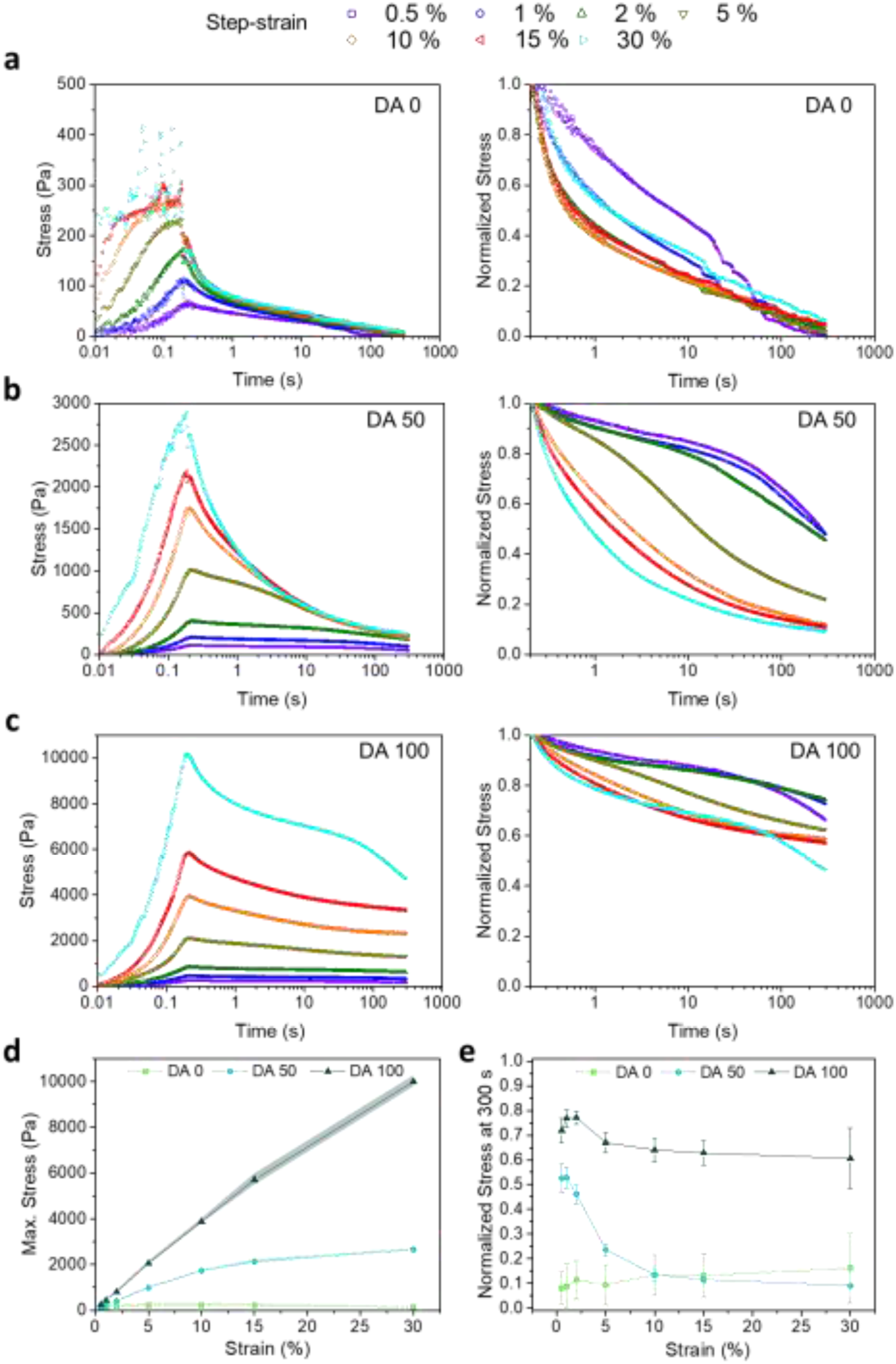
Representative step-strain stress relaxation curves as a function of increasing applied strain values from 0.5 to 30 % on **a)** DA 0, **b)** DA 50 and **c)** DA 100 hydrogels (left) with their corresponding normalized stress values (right, normalized with maximum stress values at 200 ms strain raise time); **d)** Mean maximum stress values at 200 ms (strain raise time) and **e)** Normalized stress values at the end of the stress relaxation test (300 s). Error bars indicate standard deviation, N=2 for DA 0 and DA 50 at 30 % applied strain, N=3 for all the rest.

The strain-dependent stress relaxation curves of DA X hydrogels were also dependent on the amount of X (Figure 2a). Whereas DA 0 was able to relax the induced step shear stress almost completely within 300 s at all strain values, DA 100 retained at least 60% of the stress, in agreement with its elastic nature (Figure 2e). DA 50 showed an intermediate behavior with up to 50% relaxation for shear strains <2%, whereas it relaxed >70% for larger strain values. For all hydrogels, the step shear stress was dissipated faster at higher strains, when the *γ*_F_ range was reached (Figure 2e).

The comparative relaxation behavior of DA X hydrogels (for X = 0, 25, 50, 75 and 100) at 1 % strain, i.e., within the linear viscoelastic regime, is shown in Figure 3. The shape of the curves indicates that stress is dissipated by two different relaxations processes with different time scales: a fast first one (<1 s) and a slower second one (>1 min). To analyze the relaxation mechanisms behind the two processes, the stress relaxation curves were fitted to a linear combination of two stretched exponential functions **(Eq. 1)**^28^. The function fitted well with the experimental data with r^2^ > 0.99 (Figure 3c). The values of G_0_, G_e_, A, τ_1_, τ_2_, β_1_, and β_2_ and are represented in Figure 3 d-j as a function of the hydrogel composition. The two relaxation times τ_1_ (<1 s) and τ_2_ (>1 min) differed in more than two orders of magnitude and showed an opposite dependence on the degree of the covalent crosslinking of the hydrogels. τ_1_ decreased from 0.25 s to 0.01 s as X increased from 0 to 100 (Figure 3g), i.e. the fast relaxation became faster as the covalent crosslinking degree increased. The relative contribution of this relaxation mode to stress dissipation (A parameter, Figure 3f) increased with the covalent crosslinking of the hydrogel. The shape parameter β_1_, which reflects the width of the relaxation time distribution, did not show significant changes with the hydrogel composition and is around 0.3-0.4 (Figure 3g) reflecting a 2-3 decade wide relaxation distribution. The relaxation process at longer time scales became slower in hydrogels with increasing covalent crosslinking (mean τ**_2_** increased from nearly 100 s to 560 s for DA 0 to DA 100, Figure 3h) and the strength of this relaxation decreased with X. The possible mechanisms behind these relaxations are addressed in the discussion section.

**Figure 3:**
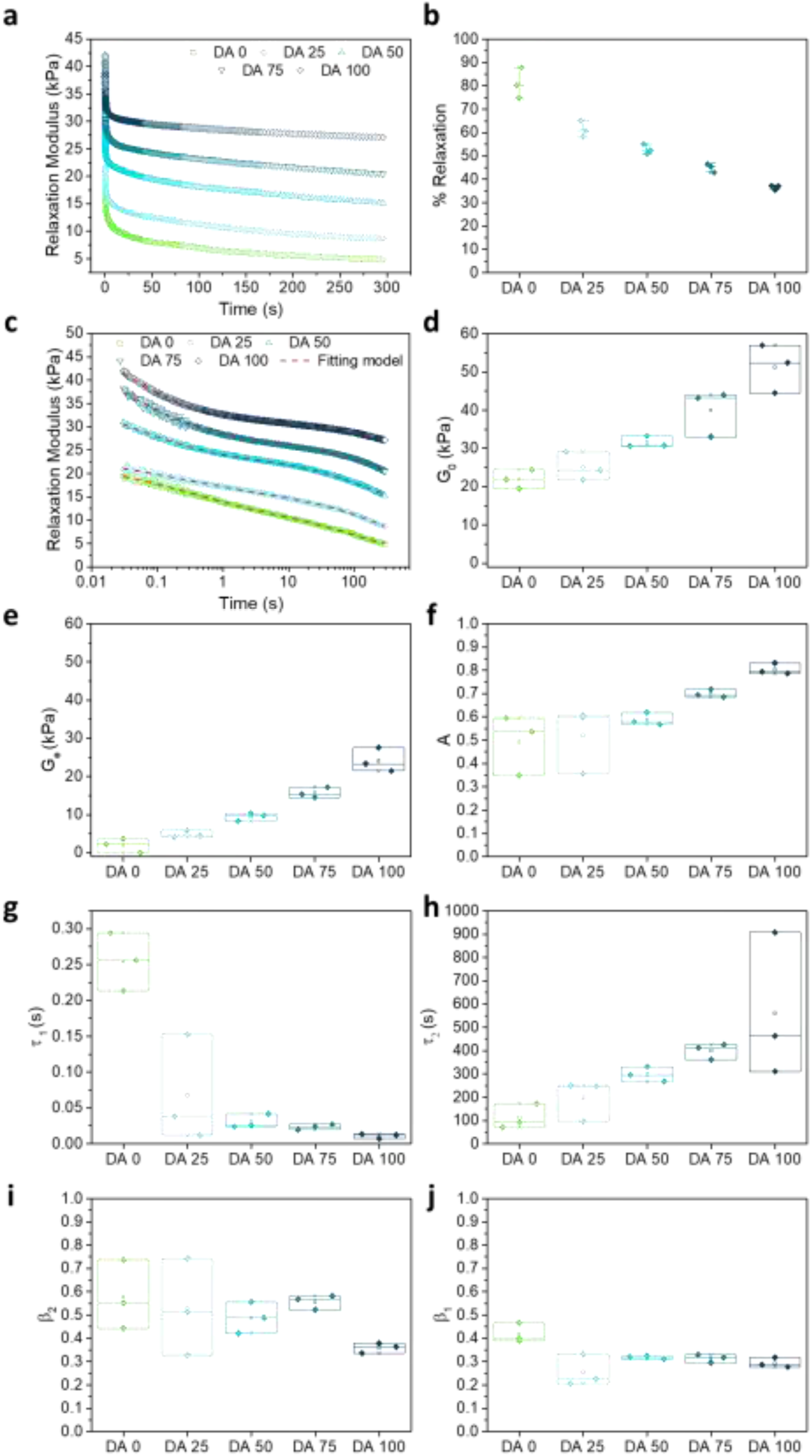
**a)** Representative stress relaxation curves of crosslinked DA X hydrogels at a constant applied strain of 1 %. **b)** Calculated normalized drop of relaxation modulus in DA X hydrogels from experiment in a) at time t = 300 s (see details in experimental section). **c)** Stress relaxation curves from a) represented on a logarithmic scale and the fits to a double stretched exponential function (red dashed lines). Data below 30 ms were not considered for the fitting since this time was required by the equipment to reach a stable strain value of 1 %; **d)** G0 values as function of X that were extrapolated linearly from stress relaxation curves in a) to time zero. **e-j)** Fitted parameters as a function of X: **e)** Ge; **f)** A (fractional contribution of relaxation 1, fast relaxation process); **g)** τ1 ; **h)** τ2 ; **i)** β1 and **j)** β2. The experimental curves of three consecutive measurements are shown in **Figure S3**. All measurements were performed at room temperature. N = 3, box represents 25 and 75 percentile values and whiskers indicate standard deviation.

### 3.5 Creep-recovery of DA X hydrogels

Figure 4a shows a creep-recovery experiment with DA X hydrogels at a constant stress of 100 Pa. In DA 25-75 hydrogels an instantaneous deformation was observed, followed by a slower deformation (creep). The instantaneous deformation was more pronounced in hydrogels with higher covalent crosslinking, while the creep process was more pronounced in hydrogels with lower PluDA concentration. DA 100 reached the plateau right after the initial deformation. DA 0 was the farthest from reaching a plateau deformation value during the 180 s of creep. When the 100 Pa stress ceased, an instantaneous and a time-dependent strain recovery was observed for all samples. DA 50-100 almost instantaneously fully recovered to their initial state, which is in agreement with their predominant elastic character as consequence of the covalent crosslinks. In comparison to that, DA25 and DA0 retained a residual deformation (Figure 4a), in agreement with their predominant physical gel character.

**Figure 4:**
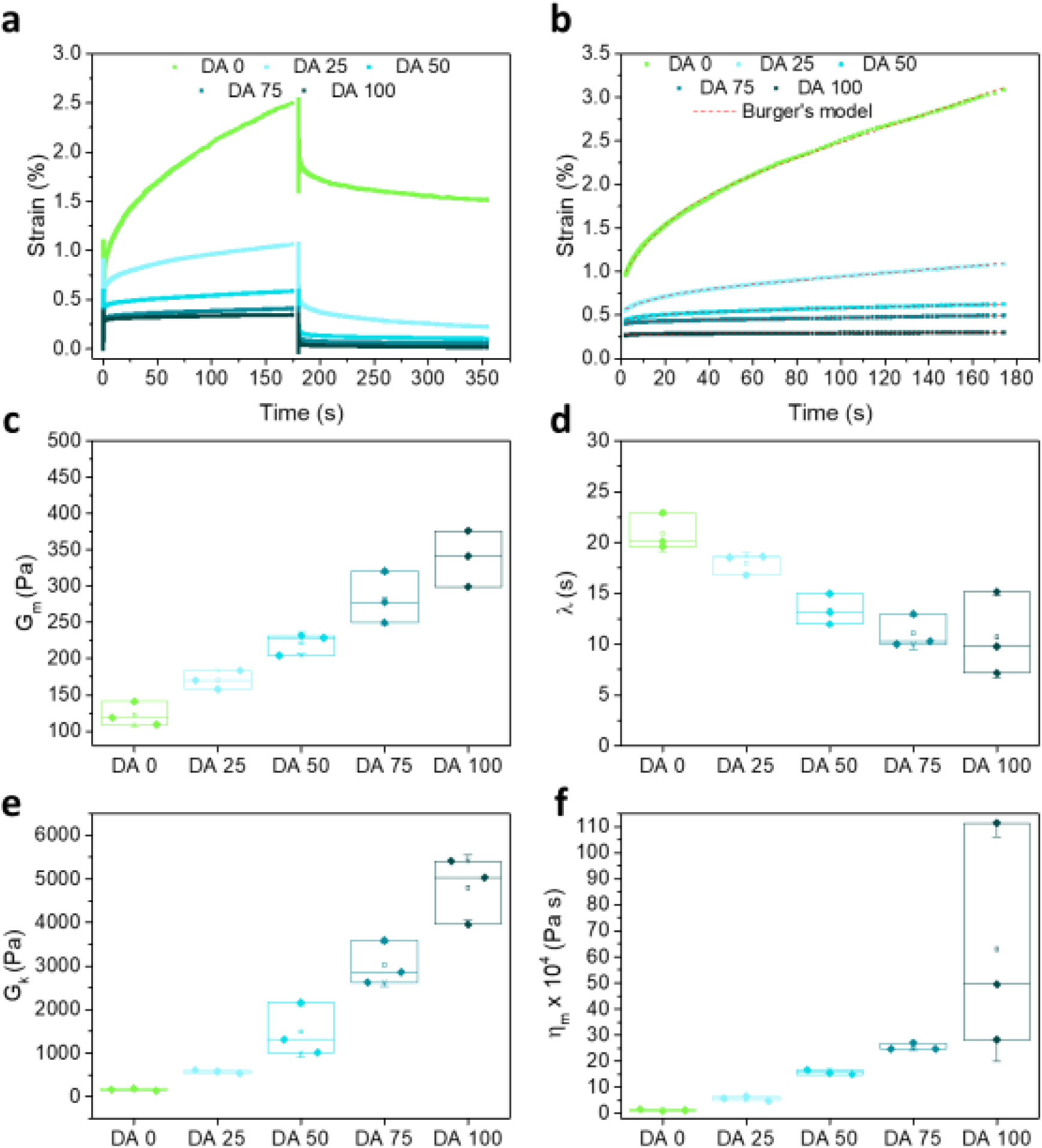
**a)** Representative creep/recovery curves of DA X hydrogels at an applied stress of 100 Pa for 180 s. The strain was monitored during stress application and also during the recovery phase for additional 180 s. **Figures S4 and S5** show all experimental curves. **b)** Creep curves and fittings to the Burgers model (**Eq. 3**, red dotted lines), **c-f)** parameters of the Burgers fitting represented as a function of X. Gm and ηm denote the spring constant of the spring and the viscosity of the dashpot in the Maxwell element, Gk is the spring constant of the dashpot in the Kelvin element and λ is the retardation time. N = 3, box represents 25 and 75 percentile values and whiskers indicate standard deviation.

The creep curves were fitted to a four-elements Burgers model (**Eq. 2**) ^30–32^ (red dashed lines in Figure 4b), consisting of Maxwell and Kelvin-Voigt elements^33,38^, and obtained fitting parameters as function of the hydrogel composition are represented in Figure 4c-f. The elastic constants G_m_ and G_k_ increased in DA X hydrogels with increasing X, and therefore an increased elasticity is observed due to a higher number of crosslinks. The reorganization of covalently linked micelles and clusters requires additional work, which causes an increment in the opposition to deformation. This higher resistance capacity contributes to elasticity. In the Burgers model, the retardation time λ is connected to the viscoelastic solid material behavior. In the micellar hydrogel model described before, the viscoelastic response is expected to depend on the intermolecular forces between micelles connected by inter-micellar covalent bonds. The shorter retardation times observed with increasing X indicate that the covalent connection between the micelles accelerates the sliding of micelles and the deformation of the hydrogel. Furthermore, η_m_ increases with X and reflects the increasing dissipative resistance, and thus lesser deformation due to the viscous component. We associate this with sliding of micelles and clusters connected by physical interactions.

The recovery part of the creep experiment was fitted using a Weibull distribution (**Eq. 3**) equation that is a modification and extension of a simple exponential relaxation. The Weibull equation fitted to the experimental data is shown in Figure 5a. The delayed viscoelastic strain recovery γ_k_ decreased with increasing covalent crosslinking in DA X hydrogels, reflecting that the presence of permanent crosslinks hinders viscoelastic deformation. The permanent irreversible strain γ_p_ was 10 times higher for DA 0 than for the other hydrogels as consequence of the absence of covalent bonds that provide elasticity.

**Figure 5:**
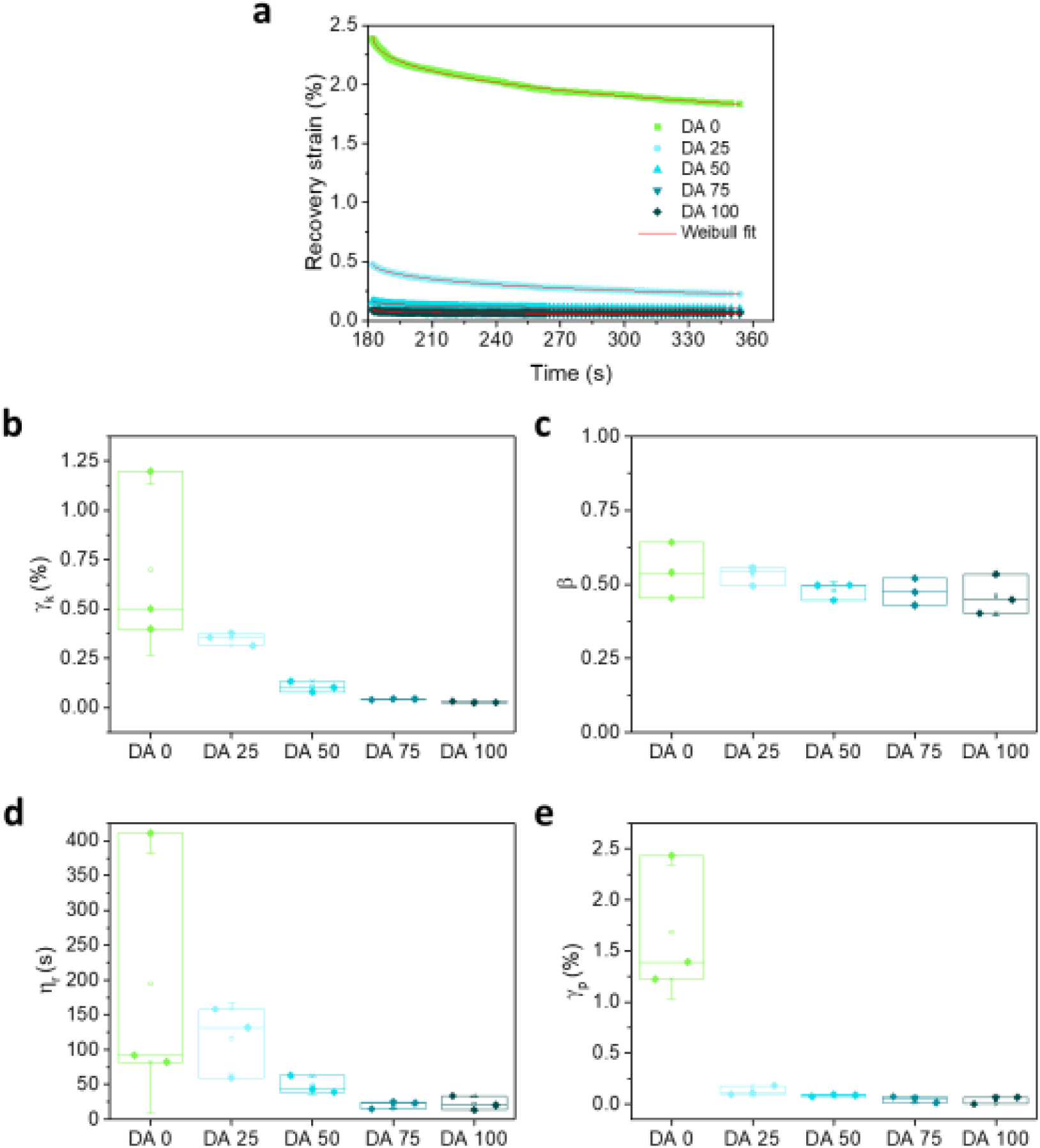
**a)** Recovery curves including fittings to the Weibull distribution function and **b-e)** parameters of the Weibull fitting as function of DAX composition. N = 3, box represents 25 and 75 percentile values and whiskers indicate standard deviation.

The elastic recovery corresponds to the instantaneous recovered deformation, that is, the Maxwell spring. γ_maxwell spring_ was calculated as:

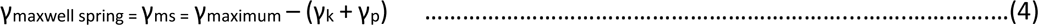

where γ_maximum_ is the strain at the end of the creep test. The elastic as well as the viscoelastic and plastic contributions to the deformation extracted from the Weibull model are represented in Figure 6.^30^ At an applied stress of 100 Pa, hydrogels DA 25-100 showed increasing elastic and decreasing viscoelastic responses with increasing covalent crosslinking, and a small plastic response with little dependence on X. DA 0 hydrogels showed a strong plastic response and a comparatively much lower elastic and viscoelastic contributions.

**Figure 6:**
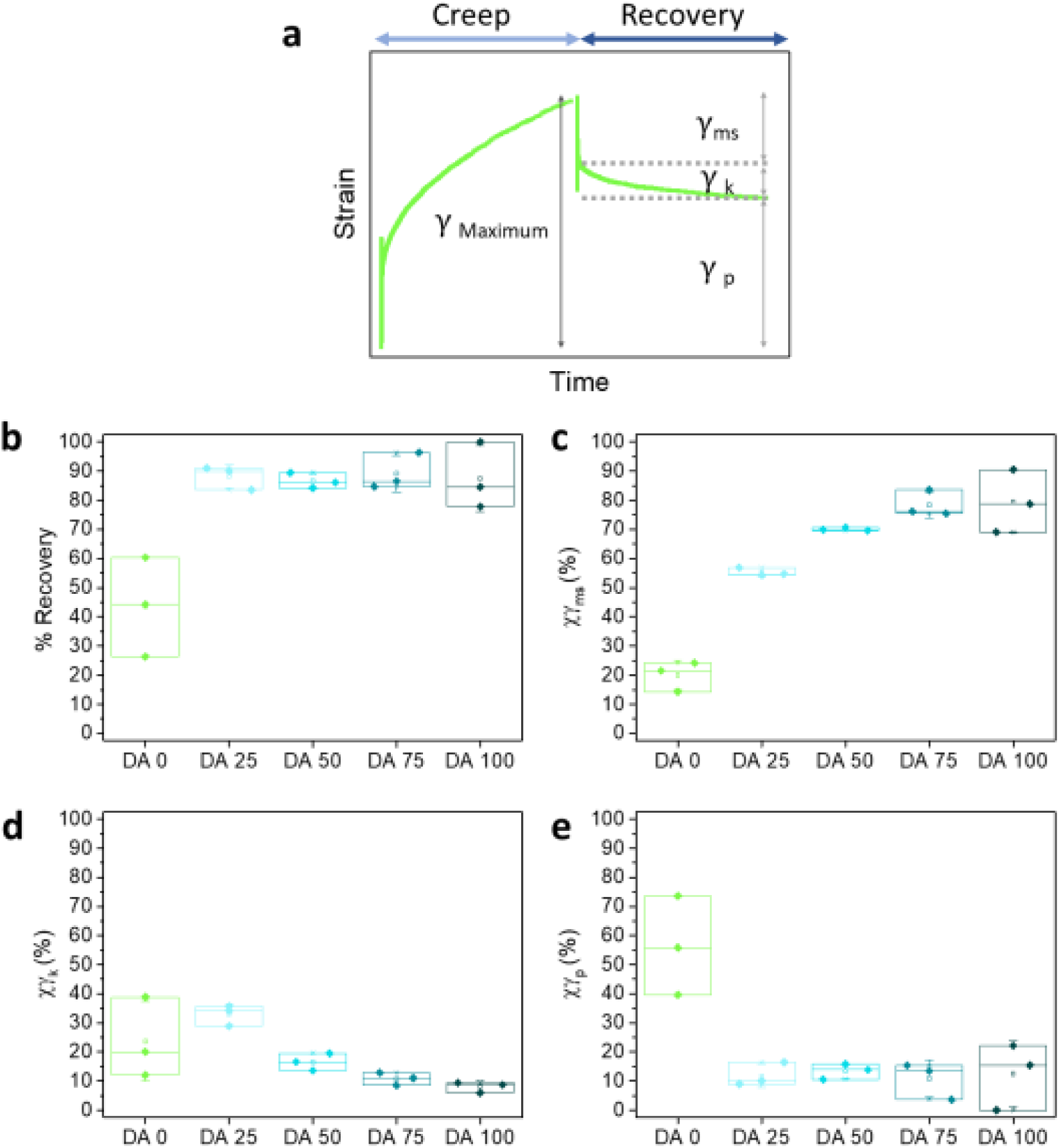
**a)** Elastic, viscoelastic and plastic contributions obtained from the creep-recovery experiment. The curve corresponds to DA 0 at an applied stress of 100 Pa; **b)** Recovery (%) calculated from recovery curves at 180 s after 100 Pa applied stress; **c-e)** Contributions of elastic, **d)** viscoelastic and **e)** plastic deformation. N = 3, box represents 25 and 75 percentile values and whiskers indicate standard deviation.

## 4 Discussion

The viscoelastic properties of 30 wt.% DA X hydrogels can be regulated by adjusting the PluDA fraction in the mixture towards hydrogels of higher elasticity and critical yield strain. The micellar DA X hydrogels are considered granular gels.^39–41^ Our rheological observations agree with this picture. In general, the internal structure of granular gels depends on the particle volume fraction and the strength of inter-particle interactions.^42^ At high particle volume fractions and weak interaction forces, gelation occurs as a consequence of the dynamical arrest of a phase separation process.^43^ In the particle-rich domains, particles aggregate and form clusters that grow until their dynamics is arrested within the 3D network.^44^ Clusters act as rigid, load bearing units and have been postulated to be the origin of elasticity in colloidal gel networks. The reversibility of the interactions between particles allows for shear-induced rearrangement of the aggregates and particles by breaking reversible bonds. As a consequence, granular hydrogels show shear thinning behavior. The observed behavior under shear of physical Pluronic hydrogels is in agreement with this picture. The inter-micellar covalent bonds introduced after physical gelation in DA X hydrogels reinforce the stability of the clusters and contribute to a stronger elastic character of the material.^41^

Previous literature discussed the behavior of Pluronic F127 hydrogels under shear.^15,45^ Pluronic micelles are considered hard spheres and their regular aggregates can undergo a first order phase transitions (analog melting, recrystallization) or reorientation as function of shear conditions (shear stress, rate or direction).^15,45^ Inter-micellar covalent bonds in DA X hydrogels internally stabilize micellar clusters and substantially affect such rearrangement mechanisms and underlying kinetics. We expect that the time scale of these rearrangements relates to the observed experimental time scales for the observed stress relaxation. Our fitting method for the stress relaxation curves of DA X hydrogels indicated two different relaxation processes with relaxation times τ_1_ (<1 s) and τ_2_ (>1 min) that depend on DA content. The fast relaxation process became faster and more prominent with increasing covalent crosslinking. We associate this relaxation to local, structural rearrangements at macroscopic, network scale which require cooperative motion of micelles.^46,47^ In contrast, the slow relaxation process could be associated to relaxation modes at microscopic scale, such as the breaking of physical bonds between micelles within clusters, which becomes slower with increasing number of covalent crosslinks.^46,48^ A recent article has studied the deformation mechanisms of 24 wt% PluDA hydrogel in an ionic liquid using rheology and SANS.^49^ The authors highlighted the possible contribution of free inter-micellar PluDA chains to elasticity by forming covalent inter-micellar bridges. Under tensile stress, the bridging chains would store mechanical energy during stretching and pull micelles back into their original positions after deformation. Although this model could explain our results, the range of stress applied was three orders of magnitude larger than the shear stress applied in our experiments and, therefore, the deformation mechanisms might differ.

## Conclusions

The presented results describe the rheological behavior of Plu/PluDA hydrogels with compositions relevant for embedding and culturing bacteria. This fundamental study can support current and future work with Pluronic-based hydrogels in the field of ELM development.

## CRediT authorship contribution statement

Shardul Bhusari: Conceptualization, Methodology, Validation, Visualization, Formal analysis, Investigation, Writing – original draft.

Maxi Hoffmann: Validation, Writing – review & editing.

Petra Herbeck-Engel: Investigation (Raman experiments).

Shrikrishnan Sankaran: Writing – review & editing.

Manfred Wilhelm: Writing – review & editing.

Aránzazu del Campo: Conceptualization, Supervision, Validation, Writing – original draft, Writing – review & editing, Funding acquisition.

## Acknowledgements

The authors acknowledge support from the Deutsche Forschungsgemeinschaft (Collaborative Research Centre, SFB 1027) and the Leibniz Science Campus on Living Therapeutic Materials, LifeMat. The authors thank Lea Fischer for useful feedback.

## Data availability statement

The raw/processed data required to reproduce these findings cannot be shared at this time due to technical or time limitations.

