## SUPPLEMENTARY INFORMATION for "Rheological behavior of Pluronic/Pluronic diacrylate hydrogels used for bacteria encapsulation"

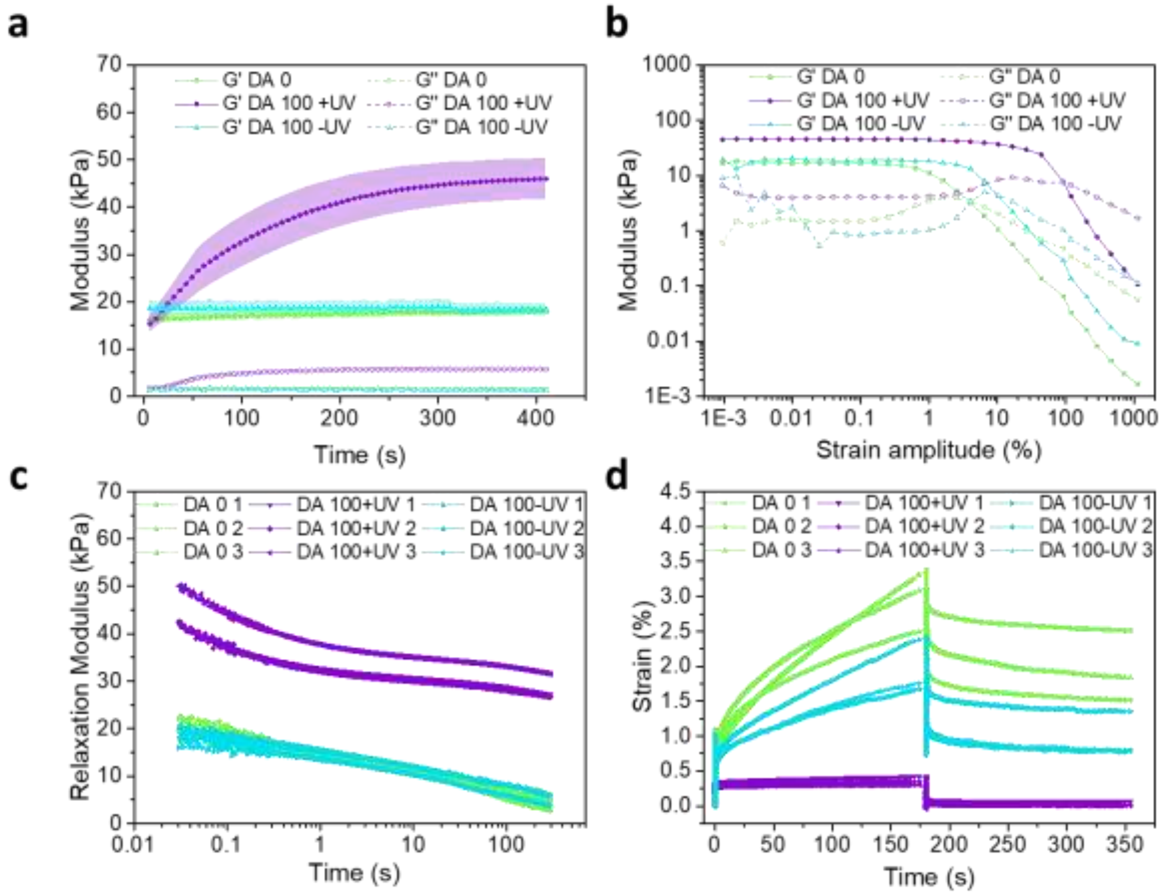


**Figure S1:** Rheological properties of DA 0 and DA 100 hydrogels before and after UV exposure and polymerization **a)** Time evolution of the G′ and G′′ (shaded regions around the data points indicate mean ± standard deviation) at room temperature; **b)** Representative strain amplitude sweep of DA X hydrogels measured at a frequency of 1 Hz; **c)** Creep/recovery curves at an applied stress of 100 Pa (Stress was applied for 180 s and then removed to measure the recovery over the next 180 s), and **d)** Stress relaxation curves at 1 % strain. In **c)** and **d)** data from 3 independent experiments are shown to visualize the reproducibility of the measurements.

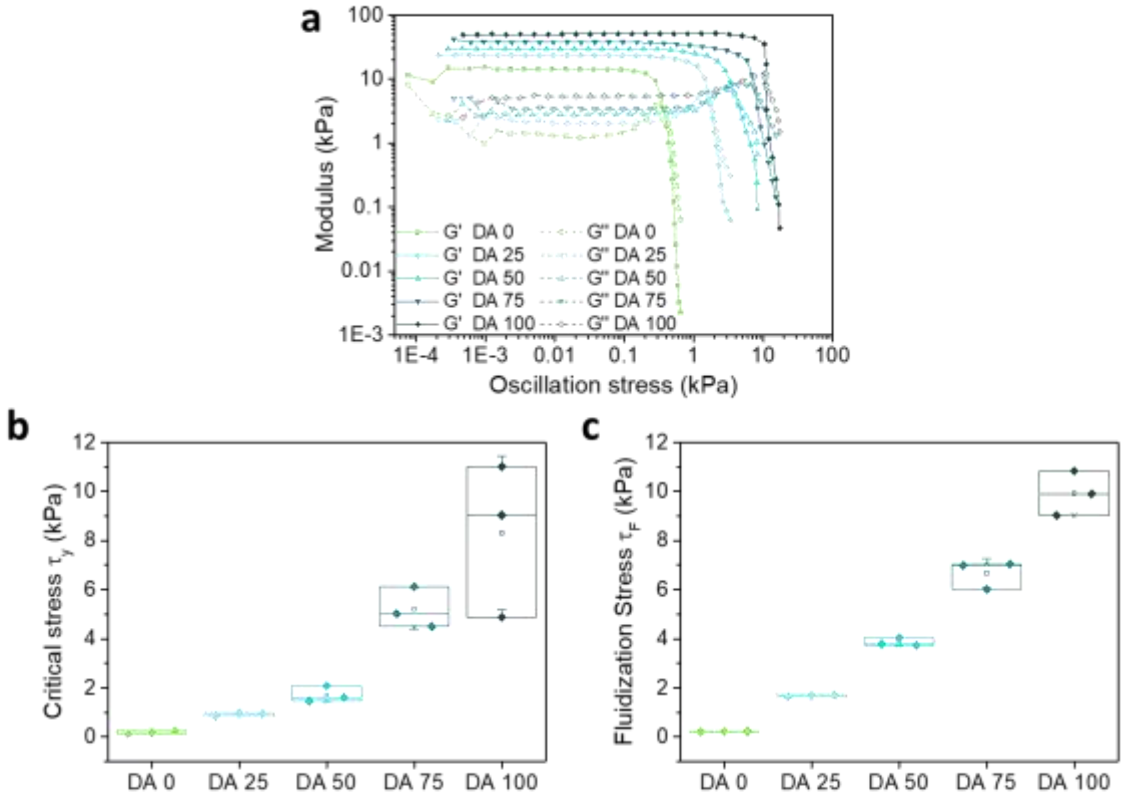


**Figure S2. a)** Representative stress sweeps and the corresponding **b)** yield stress, $\tau_y$, and **c)** stress at fluidization point, $\tau_F$ with increasing DA content in the hydrogels.

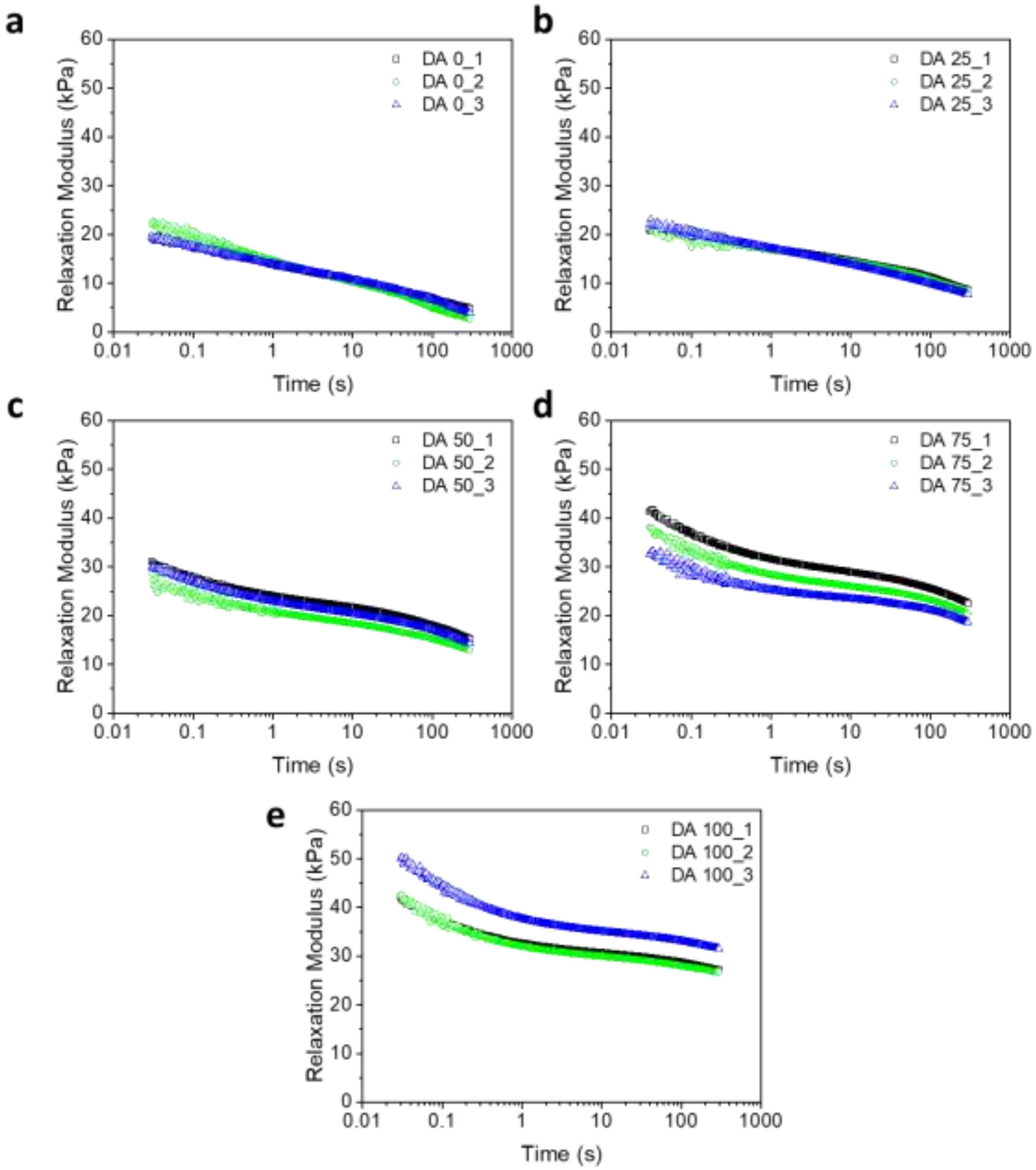


**Figure S3:** Stress relaxation curves of the DA 0-100 hydrogels at strain of 1% from three consecutive experiments used for the fitting in **Figure 3**.

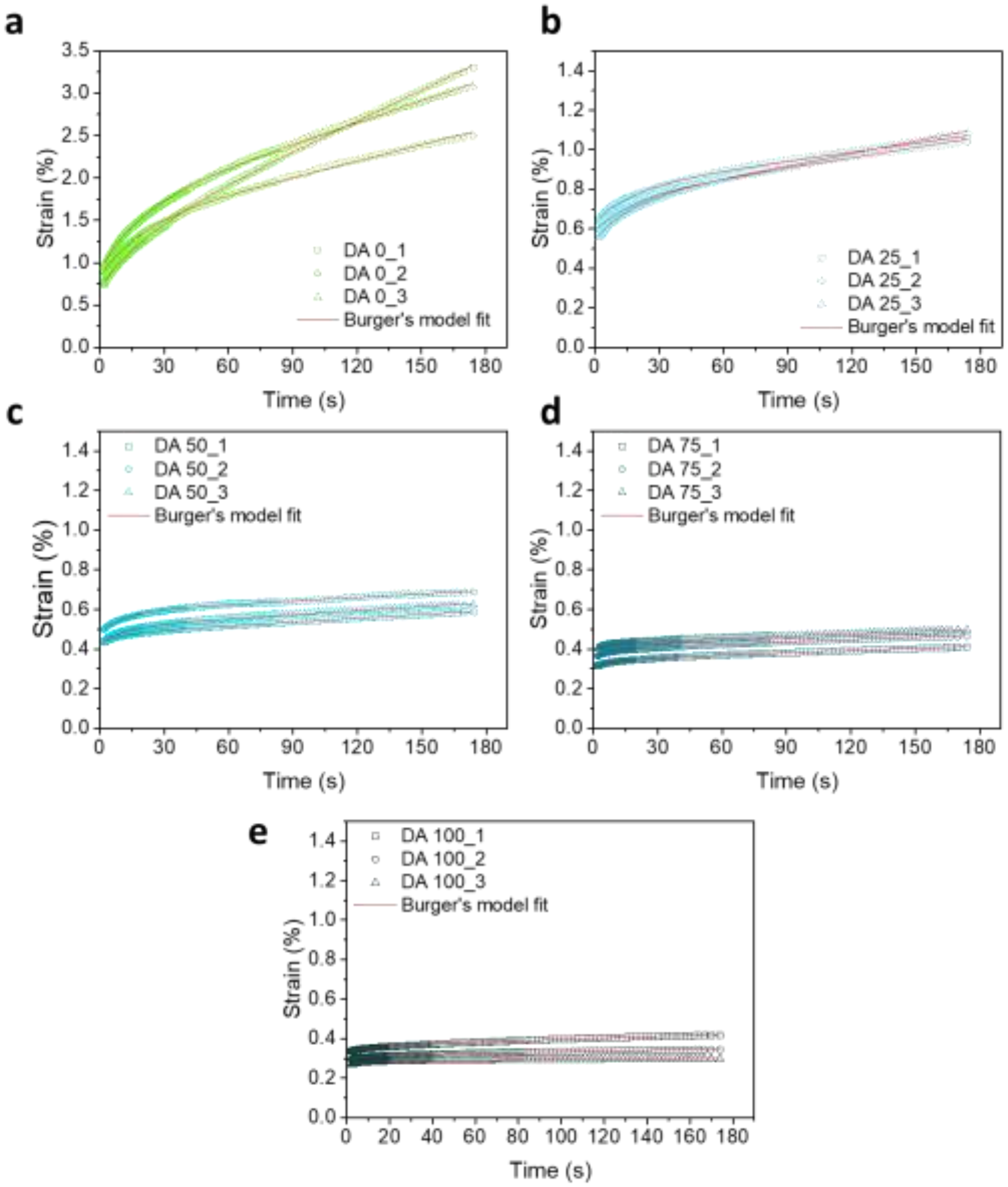


**Figure S4:** Creep strain and corresponding burgers model (**Eq. 2**) fits for the DA0-100 hydrogels.

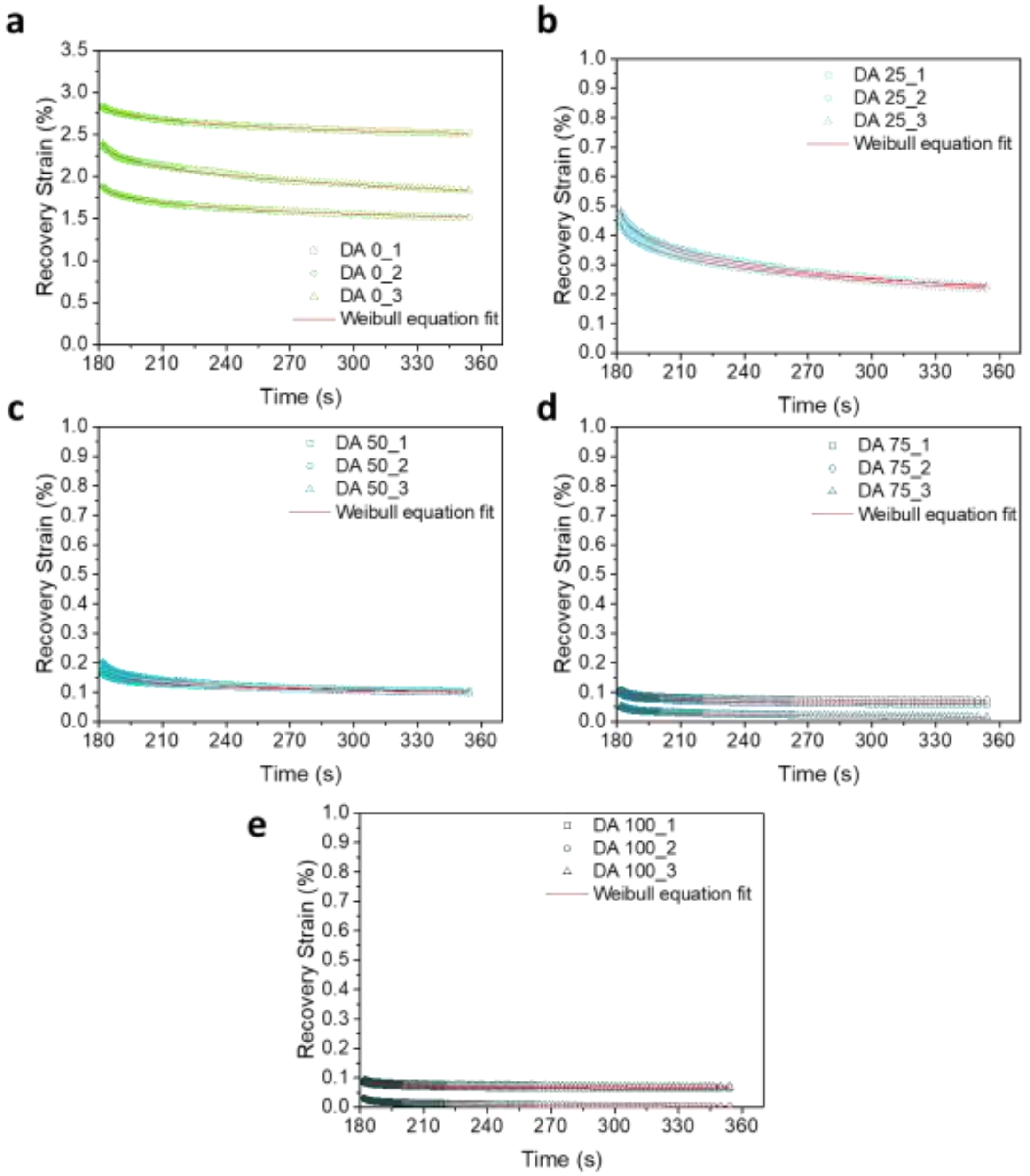


**Figure S5:** Recovery strain and corresponding fits to the Weibull equation (**Eq. 3**) for DA0-100 hydrogels.
